# UV-A/B radiation rapidly activates photoprotective mechanisms in *Chlamydomonas reinhardtii*

**DOI:** 10.1101/2020.07.21.214247

**Authors:** Ryutaro Tokutsu, Konomi Fujimura-Kamada, Tomohito Yamasaki, Keisuke Okajima, Jun Minagawa

**Author notes:** To whom correspondence should be addressed: Ryutaro Tokutsu, Division of Environmental Photobiology, National Institute for Basic Biology, 38 Nishigonaka, Myodaiji, Okazaki 444-8585, Japan, Email address, Jun Minagawa, Division of Environmental Photobiology, National Institute for Basic Biology, 38 Nishigonaka, Myodaiji, Okazaki 444-8585, Japan.

## Abstract

Conversion of light energy into chemical energy through photosynthesis in the chloroplasts of photosynthetic organisms is essential for photoautotrophic growth. However, the conversion of excess light energy into thermal energy by non-photochemical quenching (NPQ) is important for avoiding the generation of reactive oxygen species and maintaining efficient photosynthesis. In the unicellular green alga *Chlamydomonas reinhardtii*, NPQ is activated as a photoprotective mechanism through wavelength-specific light signaling pathways mediated by the phototropin (blue light) and UVR8 (ultra-violet light, UV) photoreceptors. NPQ-dependent photoprotection improves cell survival under high-light conditions; however, the biological significance of photoprotection being activated by light with different qualities remains poorly understood. Here, we demonstrate that NPQ-dependent photoprotection is activated more rapidly by UV than by visible light. We found that induction of gene expression and protein accumulation related to photoprotection was significantly faster and greater in magnitude under UV treatment compared to that under blue- or red-light treatment. Furthermore, the action spectrum of UV-dependent induction of photoprotective factors implied that Chlamydomonas sense relatively long-wavelength UV (including UV-A/B), whereas the model dicot plant *Arabidopsis thaliana* preferentially senses relatively short-wavelength UV (mainly UV-B/C) for induction of photoprotective responses. Therefore, we hypothesize that Chlamydomonas developed a UV response distinct from that of land plants.

**One-sentence summary:** In contrast to land plants, which sense short-wave UV light, the unicellular green alga Chlamydomonas senses long-wavelength UV light for photoprotective responses.

## Introduction

Light absorption is fundamental to photosynthesis, but excess light absorption damages the photosynthetic apparatus. Green photosynthetic organisms such as land plants and green algae possess light-harvesting complexes (LHC) in their photosynthetic apparatus that efficiently capture light energy (Dekker and Boekema, 2005; Minagawa and Tokutsu, 2015). This efficient light-harvesting system is advantageous under relatively weak light; however, under high-light (HL) conditions, excess light energy absorbed by LHCII can result in the formation of reactive oxygen species (Li et al., 2009), leading to photoinhibition (Takahashi and Murata, 2008). Excess light energy absorption in photosynthetic organisms is compensated for by non-photochemical quenching (NPQ), which protects photosynthesis (Horton et al., 1996; Niyogi, 1999).

NPQ is the mechanism through which excess light energy is dissipated, and is controlled by the photoprotective proteins LHC stress-related (LHCSR) and/or PSBS (Niyogi and Truong, 2013). Vascular plants lacking PSBS are deficient in NPQ activation under HL conditions (Li et al., 2000). LHCSR and PSBS also produce NPQ in the moss *Physcomitrium* (*Physcomitrella*) *patens*; mutants lacking either protein exhibit reduced rates of energy dissipation (Alboresi et al., 2010). Similarly, the unicellular green alga *Chlamydomonas reinhardtii* possesses PSBS and two LHCSRs, namely LHCSR1 and LHCSR3 (Niyogi and Truong, 2013). Chlamydomonas mutants lacking either LHCSR1 or LHCSR3 cannot survive under HL due to insufficient activation of NPQ (Peers et al., 2009; Allorent et al., 2016). LHCSR proteins in *P. patens* and Chlamydomonas function as energy quenchers (Bonente et al., 2011; Tokutsu and Minagawa, 2013; Dinc et al., 2016; Kondo et al., 2017) and/or energy distributors among photosystems (Kosuge et al., 2018).

LHCSRs in Chlamydomonas are light inducible (Peers et al., 2009; Maruyama et al., 2014). The blue-light receptor phototropin is essential for effective *LHCSR3* gene expression and protein accumulation under HL (Petroutsos et al., 2016). Moreover, the ultra-violet (UV) light receptor UVR8 can initiate UV-dependent expression of LHCSR1 and PSBS genes and proteins (Allorent et al., 2016; Tokutsu et al., 2019a). Considering that both LHCSR1 and LHCSR3 are associated with NPQ (Peers et al., 2009; Allorent et al., 2016), UV-induced activation of LHCSR1-dependent NPQ might have distinct significance compared with blue light-induced activation of LHCSR3-dependent NPQ.

Although UV and blue light are both clearly involved in the expression of photoprotective factors, the biological significance of different light wavelengths inducing different photoprotective factors in Chlamydomonas remains poorly understood. Pre-acclimation to UV enables Chlamydomonas survival of subsequent HL treatment, whereas cells not previously exposed to UV are severely bleached following HL treatment (Allorent et al., 2016; Tilbrook et al., 2016). This implies that UV-dependent activation of photoprotection in Chlamydomonas functions as “preemptive photoacclimation” before the “subsequent photoacclimation” enabled by HL-dependent photoprotection.

To further evaluate this hypothesis, we characterized the molecular and physiological responses associated with UV- and visible light-dependent activation of photoprotection. It is difficult to predict the advantages of UV-dependent activation of photoprotection in nature because 16 h of UV pre-acclimation is necessary for survival of subsequent HL treatment (Allorent et al., 2016). Under different monochromatic light conditions, however, experimental Chlamydomonas strains showed clear differences in both gene expression and NPQ activation kinetics. UV-dependent photoprotection was activated significantly faster than photoprotection activated by light with other qualities. Further analysis revealed that UV-dependent photoprotection is indispensable for HL tolerance in strains lacking LHCSR3, the photoprotective factor activated predominantly via blue-light perception.

## Results and Discussion

Although photoprotection in Chlamydomonas appears to be activated via blue-light phototropin and UV-UVR8 signaling (Allorent et al., 2016; Petroutsos et al., 2016), the precise wavelengths triggering gene expression associated with photoprotection remain unclear. To investigate this, we first analyzed LHCSR1 and LHCSR3 protein accumulation in wild-type (WT) and *npq4*-mutant Chlamydomonas strains, the latter lacking LHCSR3 (Peers et al., 2009), grown under 300–725 nm strong monochromatic light (100 μmol photons/m^2^/s) applied using an Okazaki Large Spectrograph (OLS) (Watanabe et al., 1982). After 4 h of illumination with monochromatic light, subsequent immunoblot analysis of WT cells showed distinct profiles for LHCSR1 and LHCSR3 proteins (Fig. 1). As expected, high-level LHCSR1 accumulation was observed in cells grown under light in the UV region (325–350 nm), whereas LHCSR3 accumulation was observed in response to growth under UV-A/blue (375–500 nm) and red (625–675 nm) light. UV-A-specific LHCSR1 accumulation was also observed in the *npq4* mutant, confirming that the LHCSR protein signals observed reflected accumulation of LHCSR1 rather than LHCSR3.

**Fig. 1.**
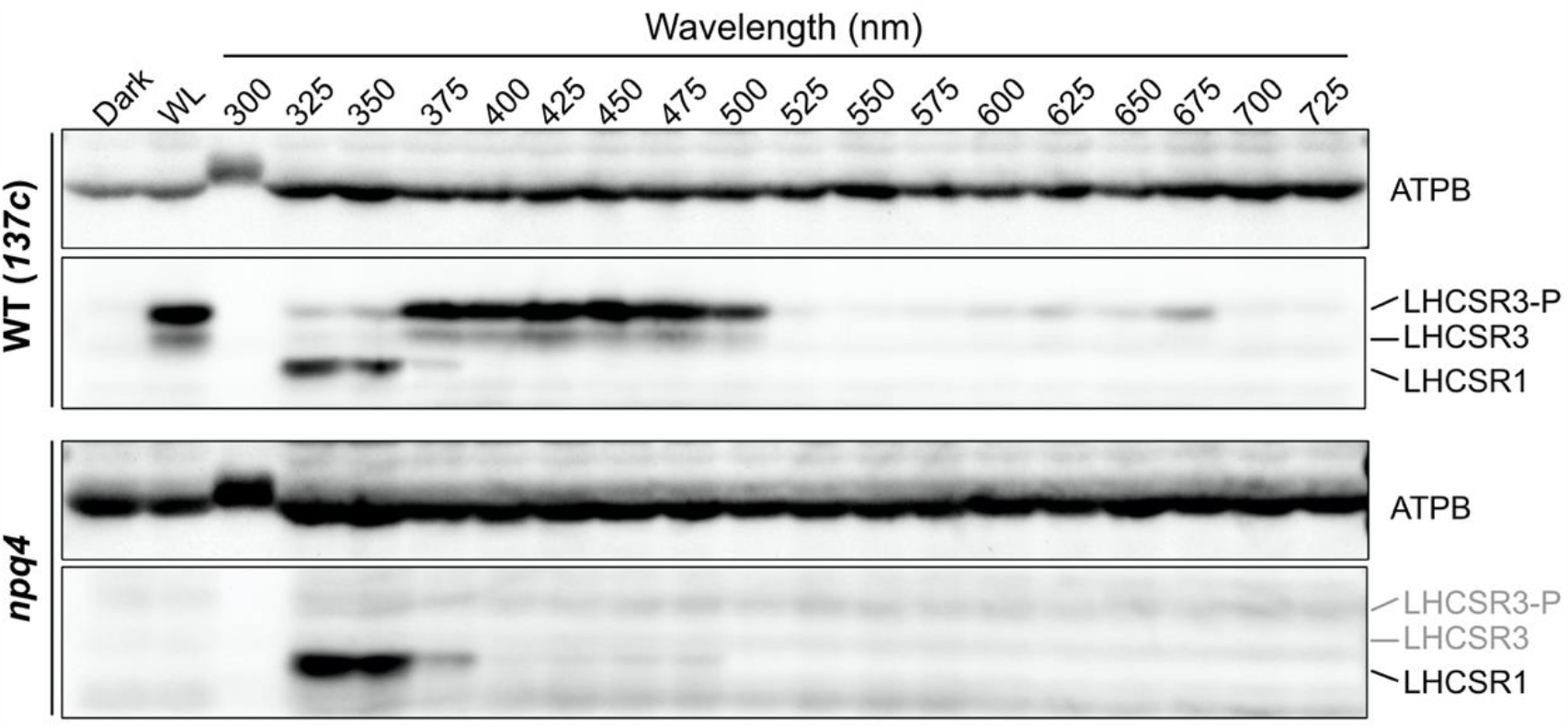
LHCSR protein levels under different wavelengths of light. Protein was extracted from samples of wild-type (WT; *137c*) and *npq4*-mutant strains. Cells were maintained in darkness (Dark) or treated with 100 μmol photons/m^2^/s of white light (WL) or different wavelengths of monochromatic light, as indicated, for 4 h. Antibodies against ATPB or LHCSRs (recognizing both LHCSR1 and LHCSR3) were used for immunoblotting analysis. Representative immunoblots from one of three replicate experiments are shown, each performed using different biological samples.

LHCSR protein accumulation was not observed in the WT or *npq4* strains under monochromatic 300-nm UV illumination. The lower level of protein accumulation under 300-nm UV also extended to the loading control protein ATPB (ATP synthase Beta subunit). Since strong UV light is known to induce photodamage of PSII via disruption of the Mn cluster in oxygen-evolving complexes (Ohnishi et al., 2005), the reduced protein accumulation observed under 300-nm UV may be attributed to cell death caused by strong photoinhibition.

To evaluate the action spectrum of LHCSR1 accumulation, we next irradiated the WT strain with monochromatic 290–350-nm UV light of relatively weak intensity (0.25 μmol photons/m^2^/s). Following 3 h of illumination, the action spectra of LHCSR1 protein accumulation exhibited peaks at 315–320 nm (Figs. 2A and S1). Although the LHCSR1 protein accumulation seemed to not be saturated under this weak UV intensity, the result was clear enough to suggest that activation of Chlamydomonas photoprotection is responsive to UV at the boundary of UV-B (280–315 nm) and in the UV-A region (315–400 nm). Consistent with the dynamics of protein levels shown in Fig. 1, LHCSR1 protein was less abundant under 300-nm UV illumination. Cells exposed to 290–300-nm UV showed similar photosynthetic activity to those exposed to other wavelengths, as indicated by the chlorophyll fluorescence parameter Fv/Fm (Fig. 2B).

**Fig. 2.**
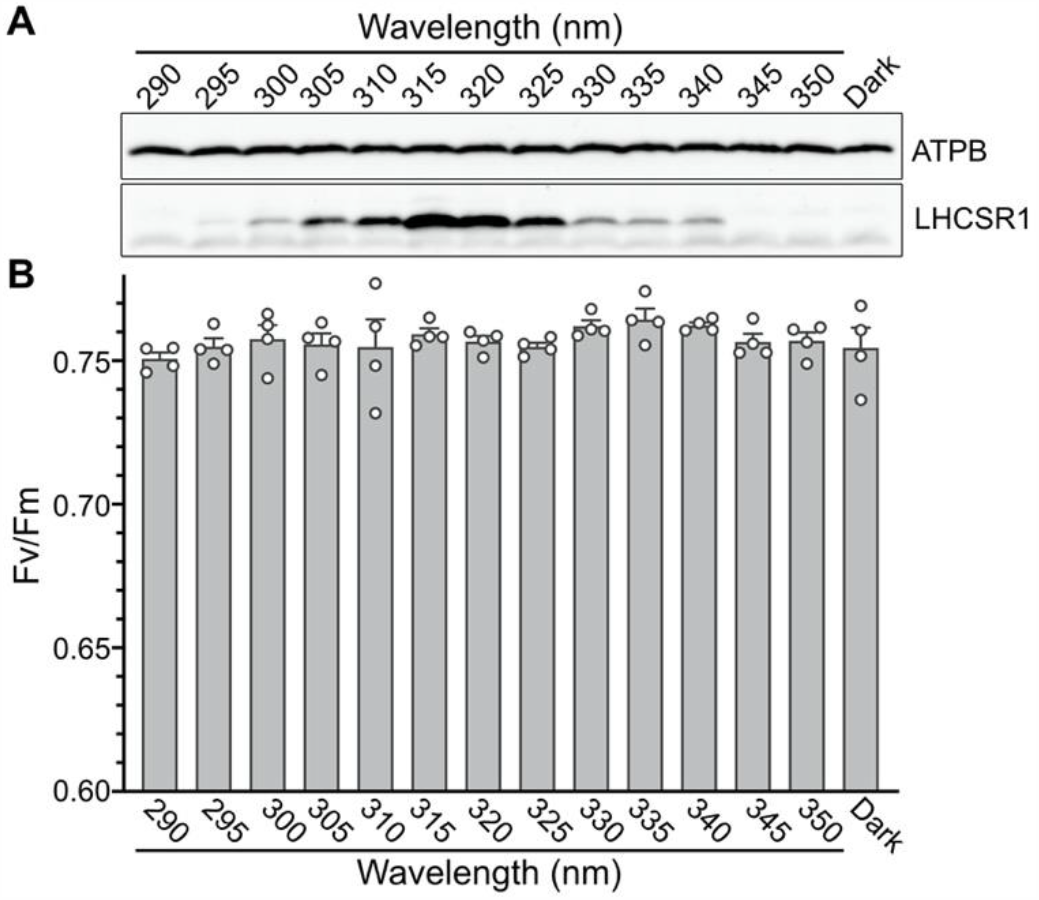
LHCSR1 protein levels and photosynthetic properties under radiation ranging from UV-A to UV-B. Wild-type cells were treated with 0.25 μmol photons/m^2^/s of monochromatic UV light of different wavelengths, as indicated, for 3 hours. A, Antibodies against ATPB or LHCSRs (recognizing both LHCSR1 and LHCSR3) were used for immunoblotting analysis. Representative immunoblots from one of three replicated experiments are shown, each performed using different biological samples. B, Photosynthetic (Fv/Fm) activities of cells from A measured using a FluorCAM system. Data are means ± SE, *n* = 4 biological replicates; raw data plots (white circles) are shown.

The action spectrum of LHCSR1 protein accumulation reported here is distinct from the UVR8 UV-absorption action spectrum in plants, which displays a peak in the UV-B and UV-C region (260–280 nm) (Brown et al., 2005; Jiang et al., 2012). This indicates that features of the UV response differ between plants and green algae, presumably as a result of differences between their habitats. Land plants are frequently exposed to harmful UV-B/C, whereas green algae are exposed mainly to UV-A because UV-B/C is rapidly quenched (absorbed) in the water (Williamson et al., 1996; Williamson and Rose, 2010). These differences suggest that green algae developed a UV response activated by UV-A instead of UV-B/C.

To determine the biological significances of UV-A/B perception in Chlamydomonas, we next investigated the activation kinetics of photoprotective responses under UV (310–330 nm), blue (470 nm), and red (660 nm) light. We first analyzed protein accumulation associated with photoprotective factors and NPQ activity. To evaluate wavelength-dependent photoprotection kinetics, the WT strain was irradiated with light with different qualities (UV, blue, and red; see Methods for details) for 240 min, and cell samples were harvested at distinct time points as indicated in Fig. 3. Subsequent immunoblot analysis again showed distinct patterns of LHCSR1 and LHCSR3 accumulation under different wavelengths (Fig. 3A). In agreement with the OLS action spectrum shown in Fig. 1, LHCSR1 protein accumulated mainly under UV light, whereas LHCSR3 accumulated under blue and red light (Fig. 3A).

**Fig. 3.**
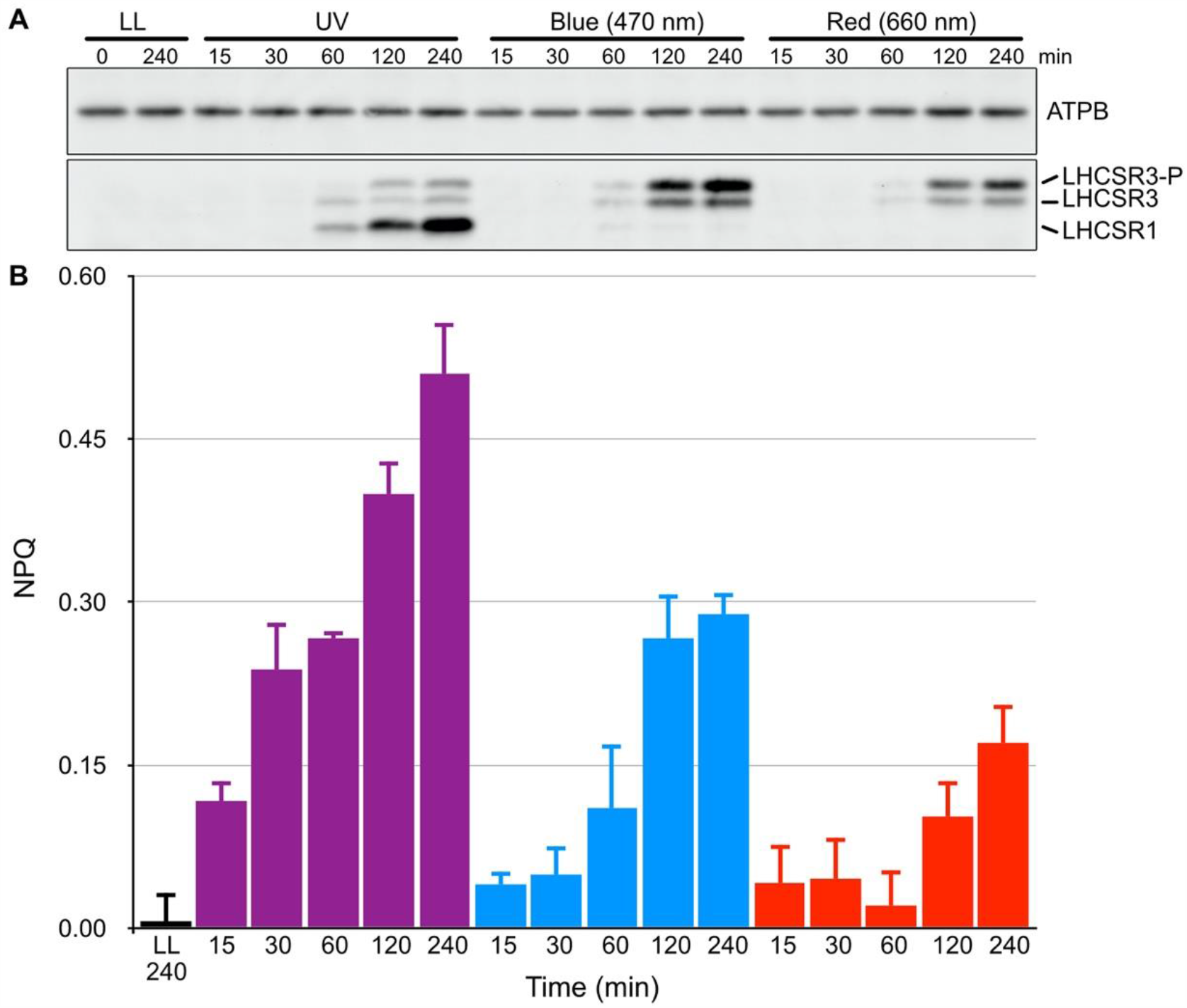
Activation kinetics of LHCSR-dependent photoprotection (NPQ) under different light conditions. A, Wild-type cell samples were collected after 15, 30, 60, 120, and 240 min of irradiation with low-level UV-supplemented fluorescent light at 10 μmol photons/m^2^/s (Fig. S3), blue (470-nm) LED light, or red (660-nm) LED light at 110 μmol photons/m^2^/s, and compared with control samples maintained under low light (LL; 30 μmol photons/m^2^/s) for the duration of the experiment. LHCSR1 and LHCSR3 protein levels were detected using an antibody against LHCSRs (recognizing both LHCSR1 and LHCSR3). The ATPB protein detected using a specific antibody was used as a loading control. Representative immunoblots from one of three replicated experiments are shown, each performed using different biological samples. B, Non-photochemical quenching (NPQ) activities of cells from A measured using a FluorCAM system. NPQ values of cells treated with LL (black bar), UV light (purple bars), blue light (blue bars), and red light (red bars) are shown. Data are means ± SE, *n* = 3 biological replicates.

Interestingly, NPQ activation kinetics were faster and NPQ was induced to a greater extent in UV-irradiated cells compared with cells irradiated with blue or red light (Fig. 3B). Although LHCSR protein accumulation was not detectable within 15–30 min of UV irradiation, NPQ activity at these time points was much higher in UV-illuminated cells than in cells treated with blue or red light. NPQ activity in cells treated with blue or red light appeared to correlate with LHCSR3 protein accumulation. These results imply that, in addition to activating LHCSR accumulation, UV treatment activates other photoprotection-related molecules such as PSBS. Considering that PSBS accumulates rapidly and temporarily before LHCSRs during activation of photoprotection (Correa-Galvis et al., 2016; Tibiletti et al., 2016; Redekop et al., 2020), it is plausible that rapid UV-dependent activation of NPQ involves both PSBS and LHCSRs in Chlamydomonas. Although it is difficult to estimate how different the kinetics of NPQ under different light are when the respective maximal values are induced by appropriate light intensity, we concluded that UV-illumination at relatively low intensity can rapidly activate NPQ when compared to blue or red light at relatively high intensity.

Since UV-A/B-dependent signal transduction is independent of photosynthesis (Allorent et al., 2016; Tokutsu et al., 2019b), it is possible that the expression kinetics of UV-A/B-induced photoprotective genes are much faster than those of genes controlled by retrograde signaling via photosynthesis under HL exposure in the visible part of the spectrum (Petroutsos et al., 2016). To further investigate whether the prompt photoprotective response of Chlamydomonas under UV light, (including rapid induction of NPQ of higher magnitude compared with that induced under red or blue light) increases with increasing photoprotection-associated gene expression, we analyzed the expression of genes encoding UV-inducible photoprotective factors (LHCSRs and PSBS) over the same time course used for the NPQ analysis shown in Fig. 3. *LHCSR1* and *PSBS1* genes were immediately induced and reached maximum expression levels after 15–30 min of UV treatment (Fig. 4, UV), while cells treated with blue or red light showed slower induction kinetics of *LHCSR3*.*2* gene expression (Fig. 4, 470 nm and 660 nm). Both *LHCSR1* and *PSBS1* genes were induced similarly by either UV or blue light to a level of at least 15 min illumination. While both genes were much more induced under UV light after 30 min of illumination, the genes’ expression was most likely regulated via UV perception. Moreover, *LHCSR1* and *PSBS1* expression was almost undetectable in red-light-treated cells (Fig. 4, 660 nm). These data imply differences in the expression kinetics of genes encoding NPQ-associated photoprotective factors induced by UV (*LHCSR1* and *PSBS1*) and those induced by visible light (*LHCSR3*.*2*) (Ballottari et al., 2016; Correa-Galvis et al., 2016; Petroutsos et al., 2016). Together with the activation kinetics of photoprotective responses shown in Fig. 3, these gene expression data indicate that UV is the most effective light quality for rapid NPQ activation in Chlamydomonas.

**Fig. 4.**
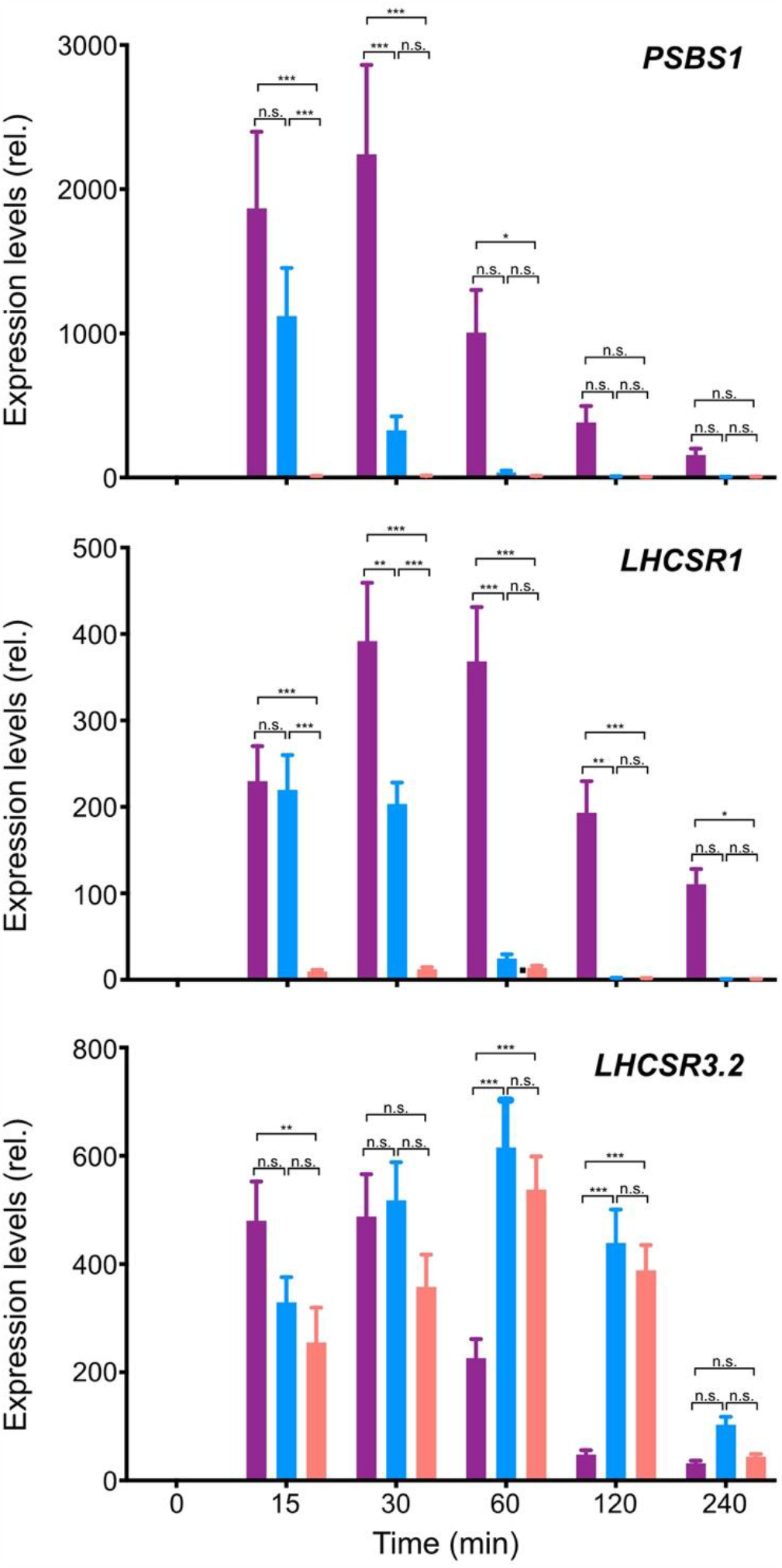
Kinetics of photoprotective gene expression under different light conditions. RNA was extracted from wild-type cell samples collected after 0, 15, 30, 60, 120, and 240 min of irradiation with low-level UV-supplemented fluorescent light at 10 μmol photons/m^2^/s (Fig. S3), blue (470-nm) LED light, or red (660-nm) LED light at 110 μmol photons/m^2^/s, as in Fig. 3. Expression levels of the photoprotection-related *PSBS1, LHCSR1*, and *LHCSR3*.*2* genes were analyzed using quantitative RT-PCR. The color bars represent expression levels each gene in UV light-(purple bar), blue light-(blue bar), and red light-(red bar) treated cells. The *CBLP* housekeeping gene was used as a control. Data are means ± SE, *n* = 3 biological replicates. Statistical significance was analyzed with multiple *t*-tests using Benjamini, Krieger and Yekutieli’s two-stage false discovery rate method procedure, with *Q* = 1%; *** denotes p<0.001; ** denotes p<0.01; *denotes p<0.05; n.s. = not significant.

The UVR8 photoreceptor is capable of perceiving UV-B in both land plants (Rizzini et al., 2011) and Chlamydomonas (Allorent et al., 2016). Using a previously obtained *uvr8* mutant (Tokutsu et al., 2019b), we performed quantitative RT-PCR analysis to evaluate whether UVR8 was responsible for UV-inducible, photoprotection-associated gene expression in Chlamydomonas. In line with a previous study (Allorent et al., 2016), the *uvr8* strain showed reduced expression of UV-induced photoprotective components, which was reflected by both mRNA and protein levels (Fig. 5). Moreover, UV-dependent rapid activation of photoprotection was severely reduced in *uvr8* cells, leading to a significantly lower NPQ light-response curve compared with the WT strain, even under low light (LL, ∼30 μmol photons/m^2^/s; Fig. 6). We also confirmed that the *uvr8* phenotypes observed could be complemented by UVR8, which was fused with the yellow fluorescent protein variant Venus and a FLAG epitope tag (Venus–FLAG) and overexpressed in the *uvr8* strain (Figs 6 and S2). These results further confirm that the UV-B photoreceptor UVR8 is responsible for the UV-dependent rapid activation of NPQ observed here.

**Fig. 5.**
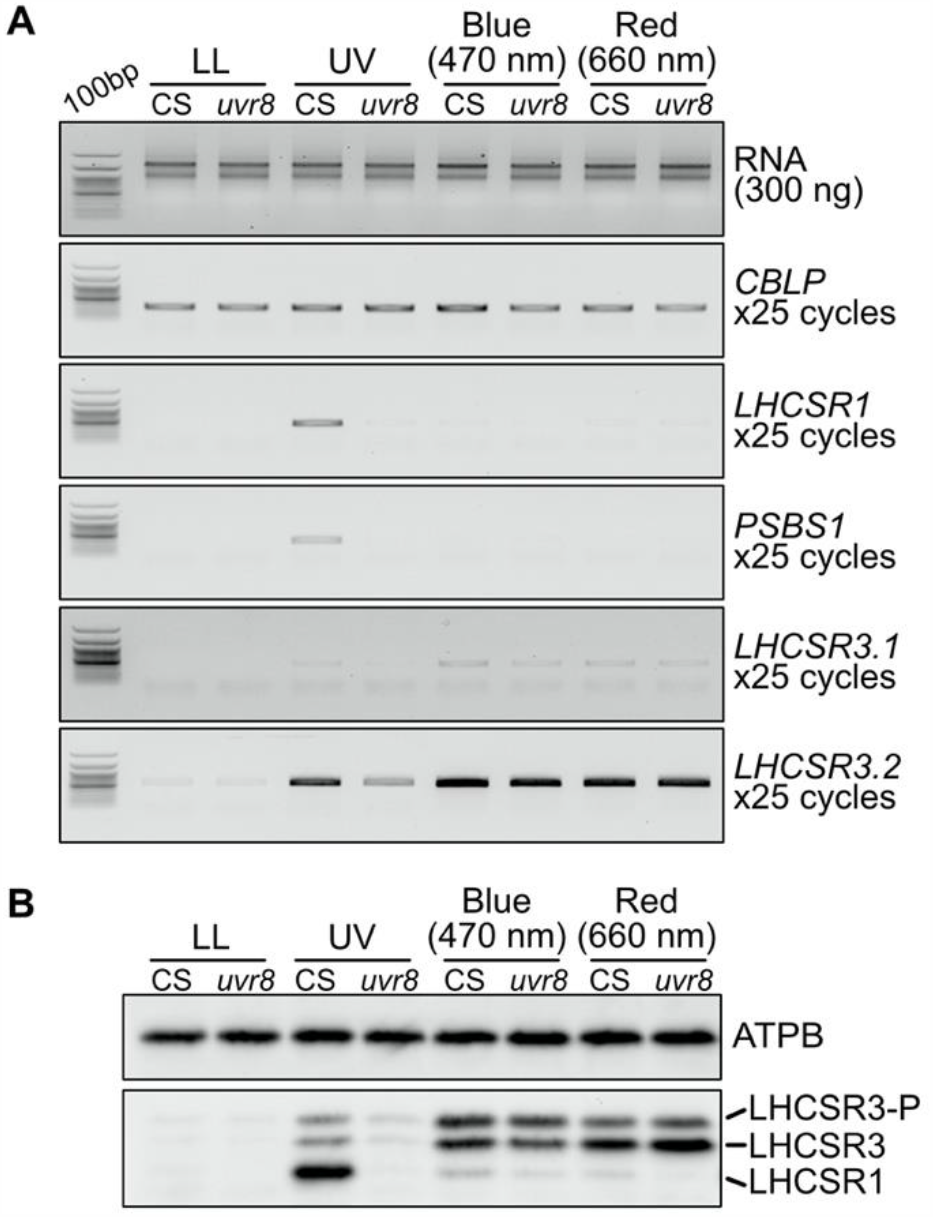
Photoprotective gene expression in the *uvr8* mutant under light of different qualities. RNA and protein were extracted from samples of control (CS; *LHCSR1–Luc717*) and *uvr8*-mutant strains illuminated for 1 h with low-level UV-supplemented fluorescent light at 10 μmol photons/m^2^/s (Fig. S3), blue (470 nm) LED light, or red (660 nm) LED light at 110 μmol photons/m^2^/s, as indicated, and compared with control samples maintained under low light (LL; ∼30 μmol photons/m^2^/s). A, Expression levels of the *PSBS1, LHCSR1*, and *LHCSR3*.*2* genes related to photoprotection were analyzed using semi-quantitative RT-PCR. The *CBLP* gene was used as a loading control. B, LHCSR1 and LHCSR3 protein levels were detected using an antibody against LHCSRs (recognizing both LHCSR1 and LHCSR3). The ATPB protein was used as a loading control. Representative gels and immunoblots from one of three replicated experiments are shown, each performed using different biological samples.

**Fig. 6.**
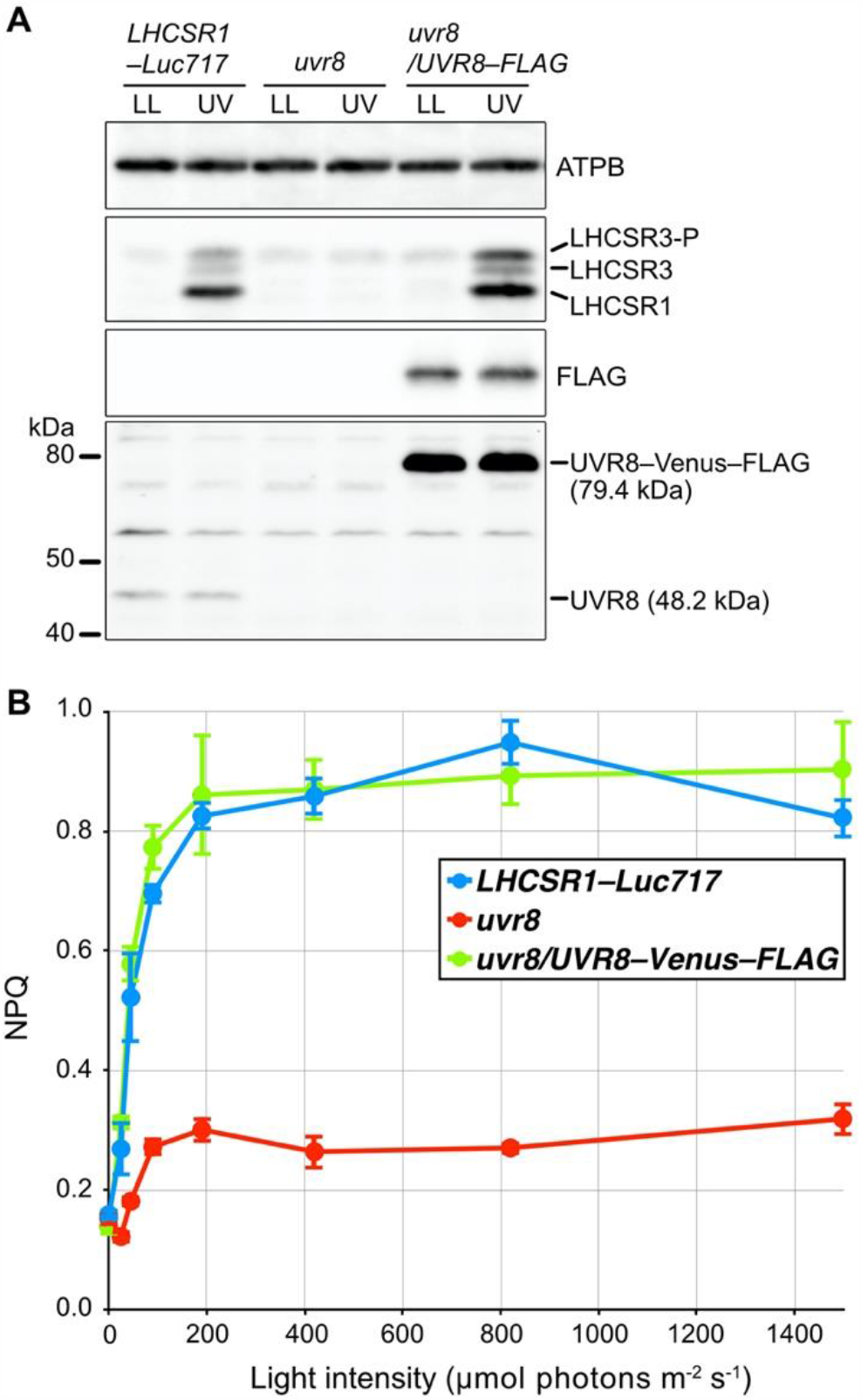
Photoprotective response of UV-treated cells. Cells of control (*LHCSR1–Luc717*), *uvr8*-mutant, and complemented (*uvr8/UVR8–FLAG*) strains were treated with low-level UV-supplemented fluorescent light (10 μmol photons/m^2^/s) for 1 h to induce photoprotective mechanisms and compared with control samples maintained under low light (LL; ∼30 μmol photons/m^2^/s). A, UVR8–Venus–FLAG and UVR8 proteins were detected using antibodies against FLAG and CrUVR8, respectively. LHCSR1 and LHCSR3 protein levels were detected using an antibody against LHCSRs (recognizing both LHCSR1 and LHCSR3). The ATPB protein was used as a loading control. B, Non-photochemical quenching was recorded using a moni-PAM system (Walz). A light-response curve was generated from measurements at 0, 45, 90, 190, 420, 820, and 1,500 μmol photons/m^2^/s. Glycolaldehyde at 10 mM final concentration was added 3 min before the measurements to interrupt the Calvin-Benson-Bassam cycle. Data are means ± SE, *n* = 3 biological replicates.

Our data establish that UVR8-dependent activation of photoprotection is established rapidly under low-level UV illumination (Fig. 6). To clarify whether this UV-dependent rapid activation of photoprotection is of biological significance under subsequent HL treatment, we next evaluated the chlorophyll bleaching phenotypes of WT and mutant Chlamydomonas strains. All strains were treated with low-level UV (10 μmol photons/m^2^/s) for 1 h to induce rapid activation of photoprotection before subsequent treatment with HL supplemented with low-level UV. Although the *uvr8* strain showed less NPQ compared with the WT (Figs 6 and S2), it did not show chlorophyll bleaching under the HL conditions applied here (Fig. 7). The photoprotection-compromised, LHCSR3-lacking *npq4* strain was also tolerant to our HL conditions and showed negligible chlorophyll bleaching (Fig. 7). However, the *npq4 uvr8* double-mutant strain exhibited significant HL-induced chlorophyll bleaching, implying that UV-dependent rapid activation of NPQ is indispensable under the HL conditions applied here in the absence of LHCSR3.

**Fig. 7.**
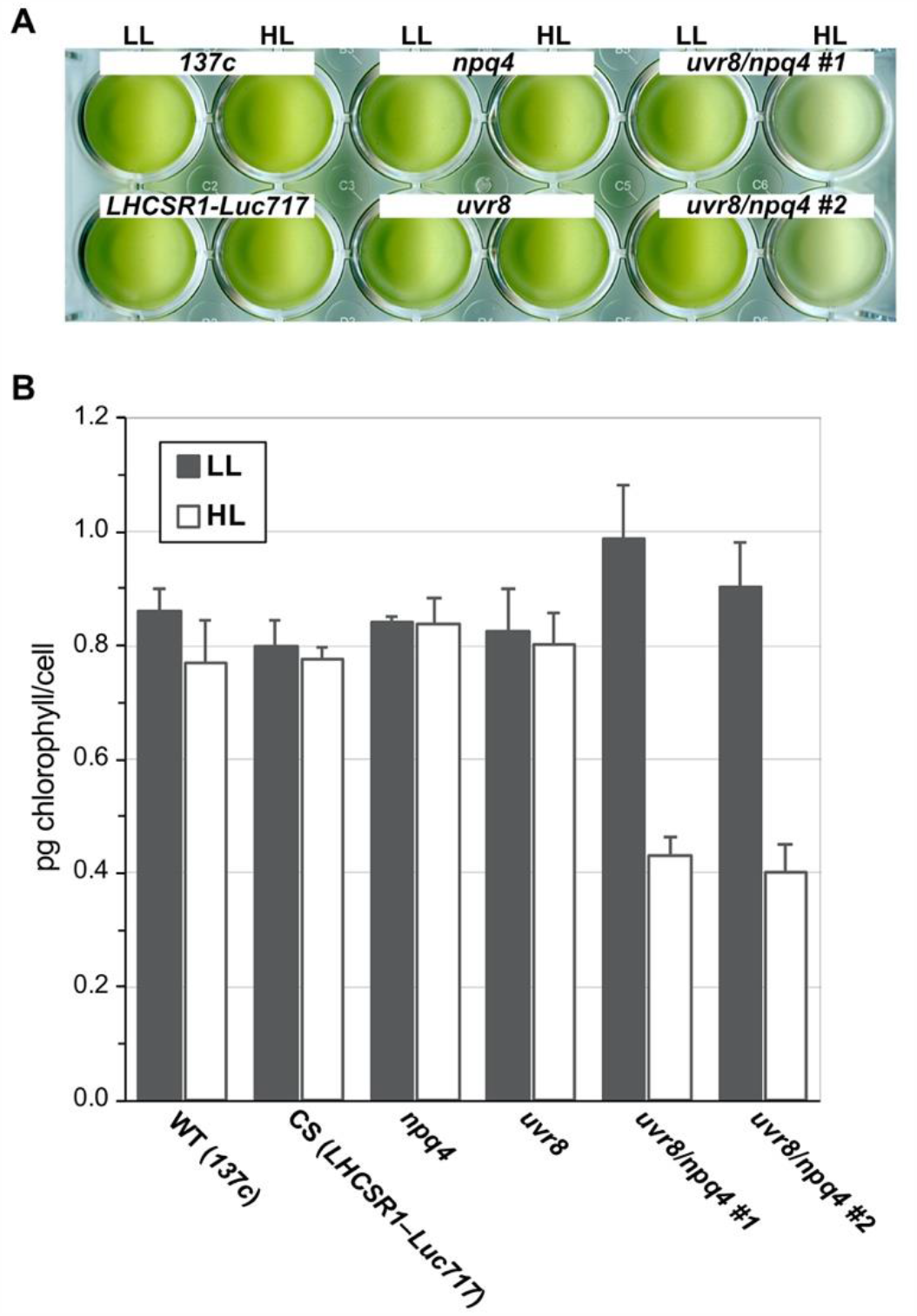
Pigment bleaching of cells induced by high light conditions. A, Wild-type (WT; *137c*), control (CS; *LHCSR1–Luc717*), and *uvr8*-, *npq4*-, and *npq4 uvr8*-mutant strains in a 24-well plate after treatment with low light (LL; 30 μmol photons/m^2^/s) or high light (HL; 500 μmol photons/m^2^/s) containing UV (UV-supplemented fluorescent light at 10 μmol photons/m^2^/s) for 10 h. Cell concentration was normalized to 5 × 10^6^ cells/mL. Representative pictures from one of three replicated experiments are shown, each performed using different biological samples. B, Chlorophyll contents of samples from A were normalized to cell number. Data are means ± SE, *n* = 3 biological replicates.

We revealed that the green alga Chlamydomonas has a distinct UV response compared with that of land plants. The Chlamydomonas UV response is initiated by relatively long-wavelength UV including that in the UV-A region (Figs 1 and S1), whereas the UV response of land plants is initiated by UV-B (Rizzini et al., 2011; Christie et al., 2012; Wu et al., 2012). It should be noted that although there is a difference in wavelength responsiveness, both Chlamydomonas and land-plant UV responses activate acclimation mechanisms that suit environmental light conditions (Brown et al., 2005; Tilbrook et al., 2016). The difference in wavelength responsiveness may be attributed to differing natural habitats. Chlamydomonas primarily inhabit the hydrosphere and wetland areas (e.g. mud, soil, wetland, and swamp), wherein the majority of UV radiation is in the UV-A region because UV-B is easily absorbed by dissolved or suspended organic/inorganic compounds in the water (Williamson and Rose, 2010). UV-A photons are much more abundant in land area, thus land plants were reported to sense UV-A via UVR8 (Rai et al., 2020). In addition to this, plants on land are exposed to more UV-B (Yin and Ulm, 2017), which causes more damage to nucleic acid and proteins compared with UV-A; therefore, land plants need to sense UV-B preferentially over long-wavelength UV. This habitat difference may explain why green photosynthetic organisms have developed a UV response to relatively long-wavelength UV (including UV-A) in the water and to relatively short-wavelength UV (mainly UV-B) in terrestrial environments. We anticipate that further insight into the UV-response strategies of these photosynthetic organisms will improve our understanding of plant evolution.

## Materials and Methods

### Algal strains and growth conditions

*Chlamydomonas reinhardtii* strain *137c* (wild type; WT) was obtained from the Chlamydomonas Center (https://www.chlamycollection.org/). The *npq4* strain was isolated in a previous study (Peers et al., 2009) and backcrossed with the WT strain several times (Kosuge et al., 2018). *DSR1* (*uvr8*) and *DSR1-comp* (*uvr8/UVR8–Venus–FLAG*) strains were generated in previous studies (Tokutsu et al., 2019a; Tokutsu et al., 2019b). The *LHCSR1–Luc717* strain harboring a reporter construct expressing a LHCSR1-Luciferase fusion (Tokutsu et al., 2019b) was used as a control strain (CS) for *DSR1*. The *npq4* and *DSR1* strains were crossed to generate double-mutant strains harboring *npq4* and *uvr8* mutations. All strains and mutants were grown in Tris-acetate-phosphate (TAP) medium (Gorman and Levine, 1965). Strains were grown under 50 μmol/m^2^/s light (Osram FL40SS D/37 white-light) at 25°C for all experiments. Once grown, cells were harvested and resuspended at 2 × 10^6^ cells/mL in high-salt (HS) medium (Sueoka, 1960), modified to include 20 mM MOPS and K_2_HPO_4_/KH_2_PO_4_ at an altered concentration of 1 mM. Cell resuspensions were then subjected to experimental light treatments as described in the text and individual figure legends.

### RT-PCR and quantitative RT-PCR

Total RNA from light-treated cells was extracted using a Maxwell RSC instrument (Promega) and a Maxwell RSC simplyRNA Tissue Kit (Promega). The RNA isolated was quantified using a QuantiFluor RNA System (Promega) prior to reverse transcription. Reverse transcription and PCR were carried out using a ReverTra Ace qPCR RT kit with gRemover (TOYOBO) and KOD FX Neo DNA polymerase (TOYOBO) in a SimpliAmp Thermal Cycler (ThermoFisher Scientific). Real-time quantitative PCR assays were performed using the KOD SYBR® qPCR Mix (TOYOBO) on the Light Cycler 96 system (Roche Diagnostics, Germany). *Δ*Ct method was used for estimating transcript abundance. For regular and quantitative RT-PCR, the housekeeping gene encoding the G protein β-subunit-like polypeptide (*CBLP*) was used as to normalize expression levels during light treatment. Primers used were as described in previous studies (Tokutsu et al., 2019a; Tokutsu et al., 2019b).

### Immunoblotting

Protein samples of whole-cell extracts (corresponding to ∼2.0 × 10^6^ cells, unless stated otherwise) were loaded onto 11% SDS-PAGE gels containing 7 M urea and blotted onto nitrocellulose membranes. Antiserum against the Beta subunit of ATP synthase (ATPB) control protein was obtained from Agrisera (AS05 085, rabbit polyclonal); antiserum against LHCSRs (recognizing both LHCSR1 and LHCSR3) was raised and affinity purified against the peptide LGLKPTDPEELK as reported previously (Tokutsu et al., 2019b); antiserum against UVR8 was raised and affinity purified against the peptide MGPDDMGTAGDSRD (Eurofins Genomics); antiserum against FLAG fusion proteins was obtained from Sigma-Aldrich (F1804, mouse monoclonal). An anti-rabbit horseradish peroxidase-conjugated antiserum (#7074, Cell Signaling Technology) or an anti-mouse horseradish peroxidase-conjugated antiserum (#330, MBL Lifescience) was used as secondary antibody. Blots were developed using EzWestLumi plus ECL detection reagent (ATTO), and images of the blots were obtained using a ChemiDocTouch System CCD imager (Bio-Rad Laboratories). The upper LHCSR3 band represents the phosphorylated form of LHCSR3 (Petroutsos et al., 2016).

### Chlorophyll fluorescence-based photosynthesis analysis

For measurement of NPQ activation kinetics (Fig. 4B), minimum (Fo) and maximum (Fm) fluorescence yield in darkness were measured using a FluorCAM (Photon System Instruments) after weak far-red (<5 μmol photons/m^2^/s) treatment for 30 min. Maximum fluorescence yield in light (Fm′) was measured following subsequent actinic irradiation at 750 μmol photons/m^2^/s for 30 s. For measurement of the light response curve (Fig. 6B), Fo and Fm were measured using a Moni-PAM system (Walz) after weak far-red (<5 μmol photons/m^2^/s) treatment for 30 min. Fm′ at different light intensities was measured following subsequent actinic irradiation at 0, 45, 90, 190, 420, 820, and 1,500 μmol photons/m^2^/s for 2 min. Glycolaldehyde, which interrupts the Calvin-Benson-Bassam cycle by inhibiting phosophoribulokinase (Takahashi and Murata, 2005), at a final concentration of 10 mM was added 3 min prior to measurements to mimic carbon limitation. Photosynthetic parameters were calculated as follows:

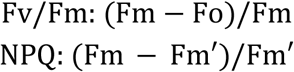

### Wavelength-dependent photoprotection assay

WT cells in HS media were irradiated with low-level UV-supplemented fluorescent light (using a ReptiSun10.0 UV fluorescent bulb (Tokutsu et al., 2019b) at 10 μmol photons/m^2^/s; Fig. S3), or 470-nm or 660-nm LED light at 110 μmol photons/m^2^/s. Total light intensities were measured using a sun spectroradiometer (S-2442 HIDAMARI mini, SOMA OPTICS, LTD.) with a range of 300 to 800 nm. Total cellular protein extracts were obtained from culture samples taken following 15, 30, 60, 120, and 240 min of light treatment.

### High-light tolerance assay

Algal strains in HS media were pretreated with low-level UV-supplemented fluorescent light (using a ReptiSun10.0 UV fluorescent bulb (Tokutsu et al., 2019b) at 10 μmol photons/m^2^/s; Fig. S3) for 1 h. Cells were then irradiated with either low light (LL; white fluorescent light at 30 μmol photons/m^2^/s) or UV-supplemented high light (HL; 500 μmol photons/m^2^/s using a ReptiSun10.0 UV fluorescent bulb at 10 μmol photons/m^2^/s; Fig. S3) for 10 h. Total cellular protein extracts were obtained from culture samples taken following 1, 2, 4, 6, and 8 h of light treatment. Chlorophyll amounts and cell numbers were determined according to the method of (Porra et al., 1989) and using a TC20 automated cell counter (Bio-Rad Laboratories), respectively. The chlorophyll amounts calculated were normalized to total cell number (pg of chlorophyll per cell). For culture photos, LL- or HL-treated strains were adjusted to 5 × 10^6^ cells/mL and transferred into a multi-well plate to be photographed.

## Author contributions

RT designed the research. RT performed transcriptional, biochemical, and pigment-bleaching analyses. RT, TY, and KF-K generated the *DSR1* mutant. RT and KO analyzed the chlorophyll fluorescence quenching of the alga. RT wrote the manuscript. JM supervised the research and provided the resources. All authors contributed to revision of the manuscript and approved the final version.

## Acknowledgements

We thank Mr. Tamaki Uchikawa and Ms. Maki Kondo for technical support in OLS experiments. Dr. Yasuhiro Kamei is thanked for fruitful discussion about UV photoreceptors. We also thank Mrs. Tamaka Kadowaki for providing technical assistance with genetic crossing of the alga. This work was supported by JSPS KAKENHI (Grant Numbers JP15H05599 and JP20H03282 to RT, and JP16H06553 to JM), the Nakajima Foundation (to RT), and NINS program for cross-disciplinary study (Grant Number 01311701 to RT). This study was carried out under the NIBB Cooperative Research for the Okazaki Large Spectrograph (16-705 and 20-609).

